# The evolution of colistin resistance increases bacterial resistance to host antimicrobial peptides and virulence

**DOI:** 10.1101/2022.02.12.480185

**Authors:** Pramod K. Jangir, Lois Ogunlana, Petra Szili, Márton Czikkely, Emily J. Stevens, Yu Yang, Qiue Yang, Yang Wang, Csaba Pál, Timothy R. Walsh, Craig MacLean

**Author notes:** Correspondence: Pramod K. Jangir (;), Craig MacLean.

## Abstract

Antimicrobial peptides (AMPs) offer a promising solution to the antibiotic resistance crisis. However, an unresolved serious concern is that the evolution of resistance to therapeutic AMPs may generate cross-resistance to host AMPs, compromising a cornerstone of the innate immune response. We systematically tested this hypothesis using globally disseminated mobile colistin resistance (MCR) that has been selected by the use of colistin in agriculture and medicine. Here we show that MCR provides a selective advantage to *E. coli* in the presence of key AMPs from humans and agricultural animals by increasing AMP resistance. Moreover, MCR promotes bacterial growth in human serum and increases virulence in a *Galleria mellonella* infection model. Our study shows how the anthropogenic use of AMPs can drive the accidental evolution of resistance to the innate immune system of humans and animals. These findings have major implications for the design and use of therapeutic AMPs and they suggest that MCR will be difficult to eradicate, even if colistin use is withdrawn.

## Introduction

The spread of antibiotic resistance in pathogenic bacteria has created an urgent need to develop novel antimicrobials to treat drug-resistant infections. Antimicrobial peptides (AMPs) are multifunctional molecules found among all kingdoms of life that act as key components of the innate immune system of metazoans by modulating immune responses and defending against invading pathogens^1,2^. AMPs are potent antimicrobials with desirable pharmacodynamic properties and a low rate of resistance evolution^3–6^. Given these benefits, there is widespread interest in the development of natural and synthetic AMPs for therapeutic use^7–9^. However, a serious concern with the therapeutic use of AMPs is that they share common physicochemical properties and mechanisms of action with AMPs of host immune system, suggesting that the evolution of bacterial resistance to therapeutic AMPs may generate cross-resistance to host AMPs^10–13^. Given that host AMPs play important roles in mediating bacterial colonization and fighting infection^14,15^, cross-resistance to host AMPs could increase pathogen transmission and virulence^16,17^.

Evolutionary microbiologists typically study the consequences of selection for antimicrobial resistance using experimental evolution. In this approach, the pleiotropic responses of bacterial populations that have been selected for increased resistance to an antimicrobial are compared with the responses of unselected control populations^18–20^. This is a powerful and tractable approach that has provided important insights into cross-resistance and collateral sensitivity, but the weakness of this approach is that the mechanisms of resistance evolution in the lab do not always match with what occurs in pathogen populations. For example, the evolution of resistance to antibiotics in many pathogens has been largely driven by the acquisition of resistance genes via horizontal gene transfer^21,22^, but conventional experimental evolution approaches focus on variation that is generated by spontaneous mutation. In this paper, we use a different approach that is based on testing the pleiotropic impacts of mobile colistin resistance genes that have become widely distributed in *E. coli* due to selection mediated by the anthropogenic use of colistin agriculture and medicine.

Colistin (polymyxin E) is an AMP produced by *Bacillus polymyxa* with similar physicochemical properties and mechanisms of action to metazoan AMPs^23,24^ (Supplementary Data 1). Colistin began to be used at a large scale in agriculture in the 1980s^25^ but it is being increasingly used as a “last-resort” antimicrobial to treat infections caused by multidrug-resistant (MDR) Gram-negative pathogens^26^. Colistin resistance has evolved in many pathogens^27–29^, but the most concerning case of colistin resistance evolution comes from mobile colistin resistance genes in *E. coli*, as exemplified by *mcr-1*^30^. Phylogenetic analyses show that *E. coli* acquired a composite transposon carrying the MCR-1 in China at some point in the 2000s^31^. MCR-1 initially spread in populations of *E. coli* from farms, where colistin was used as a growth promoter to increase the yield of chicken and pig production. However, MCR-1 became widely distributed across agricultural, human and environmental sources due to the combined effects of bacterial migration and rapid horizontal transfer of MCR-1 between plasmid replicons and host strains of *E. coli*^25,32,33^.

MCR-1 transfers phosphoethanolamine (pEtN) to lipid-A in the cell membrane, resulting in decreased net negative cell surface charge and thus lower affinity to positively charged colistin^32^. Crucially, loss of cell surface charge through membrane modification is a common resistance mechanism against cationic AMPs across bacteria^5,13^, suggesting that MCR-1 may provide cross-resistance to host AMPs. However, membrane alterations produced by MCR-1 expression are associated with clear costs^34^, and it is equally possible that membrane remodelling could generate collateral sensitivity to AMPs, as has been observed with antibiotic resistance genes^35^.

In this paper, we test the hypothesis that evolving colistin resistance via MCR genes acquisition generates selection in bacteria against host AMPs by increasing AMP resistance. MCR-1 is usually carried on conjugative plasmids from a diversity of plasmid incompatibility types (such as IncX4, IncI2, IncHI2 and IncP1) that carry a large number of housekeeping and cargo genes^31,34^. We assessed the importance of this diversity by transferring a diversity of naturally occurring plasmids and synthetic MCR-1 expression vectors to a single recipient strain of *E. coli*. To assess the impact of MCR on resistance to host AMPs, we screened a panel of strains carrying naturally occurring and synthetic MCR plasmids against a collection of AMPs. Given the importance of agricultural animals as reservoirs of MCR-1, we tested AMPs that play important roles in the innate immunity of humans, pigs and chickens (Table 1). Next, we examined the role of MCR in complex host environments and bacterial virulence using human serum resistance assays and *in vivo* virulence assays in the *Galleria mellonella (G. mellonella)* infection model system. The key innovation in this study is that we have taken a systematic approach to testing the pleiotropic effects of the dominant mechanism of colistin resistance evolution, including assessing the impact of AMP resistance on bacterial virulence.

**Table 1.** List of natural MCR plasmids and AMPs used in this study.

| MCR plasmids |  |  | AMPs |  |  |
| --- | --- | --- | --- | --- | --- |
| Name (type) | Size | *mcr* gene | Name | Abbrev | Major cell and tissue sources |
| PN16 (IncI2) | 60,488 bp | *mcr-1* | LL-37 cathelicidin | LL37 | Epithelial cells of the testis, skin, gastrointestinal tract, respiratory tract, and in leukocytes, such as monocytes, neutrophils, T cells, NK cells, and B cells |
| PN21 (IncI2) | 60,989 bp | *mcr-1* | Human beta-defensin-3 | HBD3 | Neutrophils and epithelial surfaces (e.g., skin, oral, mammary, lung, urinary, eccrine ducts, and ocular) |
| PN23 (IncX4) | 33,858 bp | *mcr-1* | Cecropin P1* | CP1 | Small intestine |
| PN42 (IncX4) | 32,995 bp | *mcr-1* | PR39 | PR39 | Mucosa and lymphatic tissue of the respiratory tract |
| WJ1 (IncHI2) | 266,119 bp | *mcr-3* | Protegrin 1 | PRO1 | Bone marrow, leukocytes and neutrophils |
| 481 (IncP1) | 53,660 bp | *mcr-3* | Prophenin-1 | PROPH | Bone marrow and leukocytes |
|  |  |  | PMAP-23 | PMAP23 | Myeloid tissue, bone marrow and liver |
|  |  |  | Chicken cathelicidin-2 | CATH2 | Bone marrow, respiratory tract, gastrointestinal tract, normal intact skin and multiple lymphoid organs |
|  |  |  | Fowlicidin 3 | FOW3 | Bone marrow, lung and spleen |
|  |  |  | Colistin | COL | - |

\*from pig intestinal parasitic nematode

## Results

### Host AMPs select for MCR-1

To assess the consequences of MCR-1 aquisitions without any confounding effects from backbone and cargo genes found in naturally occurring MCR-1 plasmids, we cloned MCR-1 and its promoter into a non-conjugative expression vector (pSEVA121) that has a similar copy number to naturally occurring MCR plasmids (approx. 5 copies per cell). As a first approach to assess the impact of MCR-1 on resistance to host AMPs, we measured the competitive ability of pSEVA:MCR-1 across a concentration gradient of a randomly selected representative set of host AMPs and colistin, which acts as a positive control for *mcr-1* selection (Figure 1).

**Figure 1.**
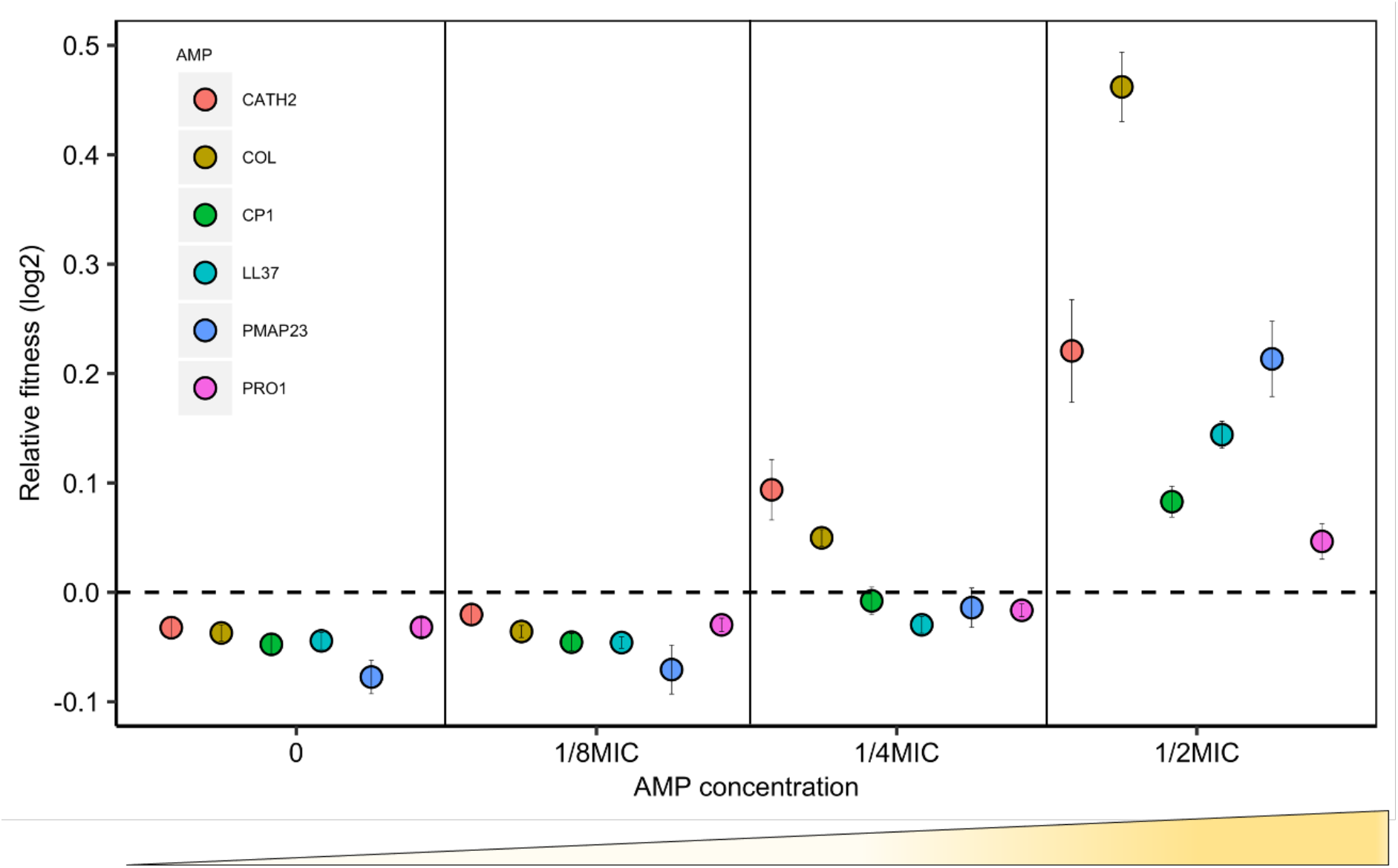
Sub-MIC doses of AMPs generate selection for MCR. *E. coli* carrying MCR-1 expression vector (pSEVA:MCR-1) or an empty vector control (pSEVA:EV) were competed against a tester strain carrying a chromosomally integrated GFP across a range of AMP concentrations (*n* = 6 biological replicates per competition). Plotted points show the competitive fitness of the MCR-1 expressing strain relative to the empty vector control (+/- s.e). To facilitate comparisons across AMPs, fitness is plotted as a function of relative AMP concentration, and the dashed line represents equal fitness.

Consistent with previous work^34^, MCR-1 imposed a significant fitness burden in the absence of AMPs, reducing competitive ability by 3% (*P* **=** 1.174e-15, two-sided Mann– Whitney U test, Supplementary Figure 1). However, MCR-1 provided a gain of a significant competitive fitness advantage at concentrations of host AMPs between ¼ and ½ of minimum inhibitory concentration (MIC) (Figure 1, Supplementary Data 2). Although MCR-1 provided a greater fitness advantage in the presence of colistin as compared to host AMPs, the minimal selective concentration for colistin, ¼ MIC, was only marginally lower (Figure 1). It is important to note that the sub-MIC concentrations required for the selection of MCR-1 overlap with the range of physiological concentration of host AMPs. For example, the concentration of LL-37 required to select for MCR-1 (∼3.4 μM) falls well within the reported physiological concentration range (up to 10 μM)^36,37^.

### MCR increases resistance to host defense AMPs

To test the hypothesis that MCR increases resistance to host AMPs more broadly, we measured the resistance of MCR-*E. coli* to a panel of AMPs. Given the importance of agricultural animals as reservoirs of MCR^30^, we tested AMPs that are known to play important roles in the innate immunity of chickens, pigs and humans. The panel of AMPs used in our assay have diverse mechanistic and physicochemical properties, and include AMPs that are known to have clinical relevance and play key roles in mediating innate immunity (Table 1 and Supplementary Data 1). For example, the human cathelicidin LL-37 and defensin HBD-3 have immunomodulatory activities in addition to their antimicrobial activity^8,38,39^.

We tested the AMP resistance of both *E. coli* carrying pSEVA:MCR-1, which provides a clean test for the effect of the *mcr* gene, and transconjugants carrying diverse MCR-1 and MCR-3 natural plasmids. These plasmids represent the dominant platforms for MCR found in clinical and agricultural sources in South East Asia^31^, and plasmid diversity may play an important role in mediating the effect of *mcr* due to variation in plasmid copy number and the effect of other plasmid genes on AMP resistance. AMP resistance genes typically give much smaller increases in resistance than antibiotic resistance genes, typically on the order of 1-2 fold increases in MIC^40^. To take the quantitative nature of AMP resistance into account, we measured MIC using an established assay that had the sensitivity to capture small differences in bacterial resistance^40^ (Figure 2).

**Figure 2.**
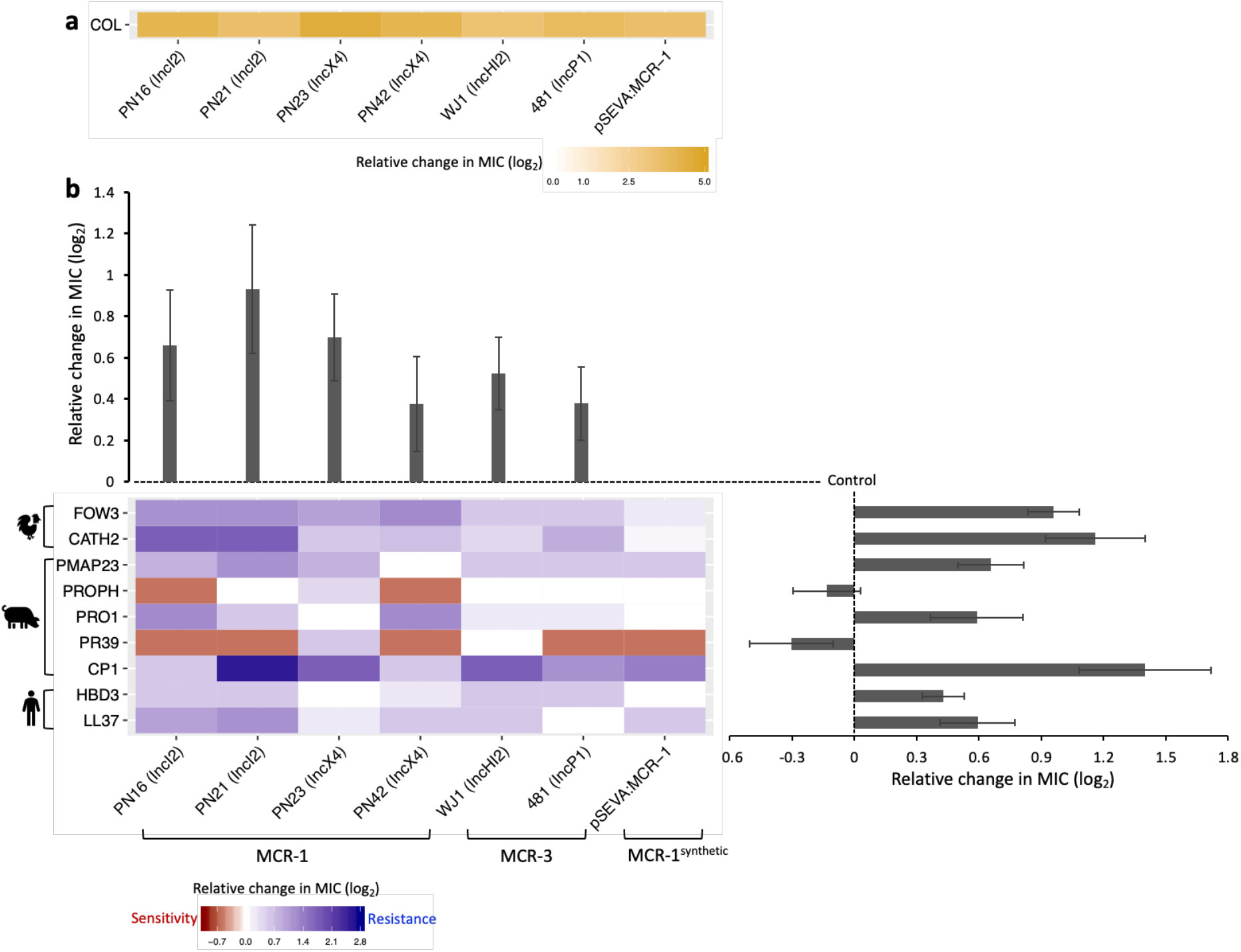
MCR-mediated changes in bacterial susceptibility to host AMPs. Heatmaps depict the effect of MCR plasmids on resistance to colistin **(a)** and host AMPs **(b)**. Bacterial susceptibility to AMPs was tested by measuring MICs and changes in resistance were assessed relative to control strains lacking MCR (*n* = 3 biological replicates per MIC). Natural plasmids carried either MCR-1 or MCR-3 are shown according to plasmid incompatibility group. Resistance for these plasmids was measured relative to the *E. coli* J53 parental strain. The impact of the synthetic pSEVA:MCR-1 plasmid on resistance was measured relative to a strain with a pSEVA empty vector. Bar plots show average changes in MIC for natural MCR plasmids, and did not include pSEVA:MCR1 (+/- s.e; *n* = 9 for host AMPs, *n* = 6 for plasmids).

On average, MCR plasmids provided increased resistance to host AMPs (mean increase in log^2^ MIC = 0.58; s.e = 0.075; t = 9.05; *P*<0.0001; Figure 2; Supplementary Data 3). However, the average change in resistance conferred by MCR plasmids varied significantly between AMPs (F_8,40_ = 7.85; *P*<0.0001), as MCR plasmids increased resistance to most AMPs, but generated collateral sensitivities to both PROPH and PR39 (Figure 2b). These AMPs have unique physicochemical properties (Supplementary Figure 2), including high proline content (Supplementary Figure 2b), which has been shown to be a common property of intracellular-targeting AMPs, as opposed to membrane-disrupting AMPs (Supplementary Data 1)^41,42^. These results agree with recent work showing that collateral sensitivity interactions are frequent between membrane-targeting and intracellular-targeting AMPs^40^.

The AMP resistance profile of MCR plasmids was comparable to pSEVA:MCR-1, suggesting that changes in resistance observed in MCR plasmids were caused by MCR, and not by other genes present on these plasmids. To further test this idea, we replaced the *mcr-1* gene on an IncX4 natural plasmid (PN23 IncX4) with an ampicillin resistance marker, which is not known to have any effect on AMP resistance. As expected, deletion of *mcr-1* gene resulted in a wild-type level of resistance to AMPs (Supplementary Figure 3). Altogether, these results suggest that the observed AMP resistance phenotype is largely due to the pleiotropic effects of MCR gene and is not distorted by other genes present on natural plasmids.

The high level of colistin resistance provided by MCR-1 plasmids (Figure 2a, mean change in log^2^MIC=3.98, range=3.5-4.5) in comparison to host AMPs is striking given the broad mechanistic and physicochemical similarities between colistin and membrane-targeting AMPs. To better understand the origins of the high colistin resistance phenotype associated with *mcr-1*, we cloned the closest known homologue of MCR-1, from the pig commensal *Moraxella* (MCR-MOR, ∼62% identity with MCR-1) into pSEVA121^32,43^. In line with previous work, MCR-MOR expression provided a small increase (5.9-fold) in colistin resistance as compared to MCR-1 (13.2-fold) ^44,45^ (Figure 3a). In contrast, MCR-MOR expression did not increase resistance to host AMPs tested with the exception of CP1, which is secreted in the pig gut (Figure 3b-d). The increased CP1 resistance conferred by MCR-MOR is intriguing, and it suggests that this AMP may impose a significant selective pressure in the pig gut.

**Figure 3.**
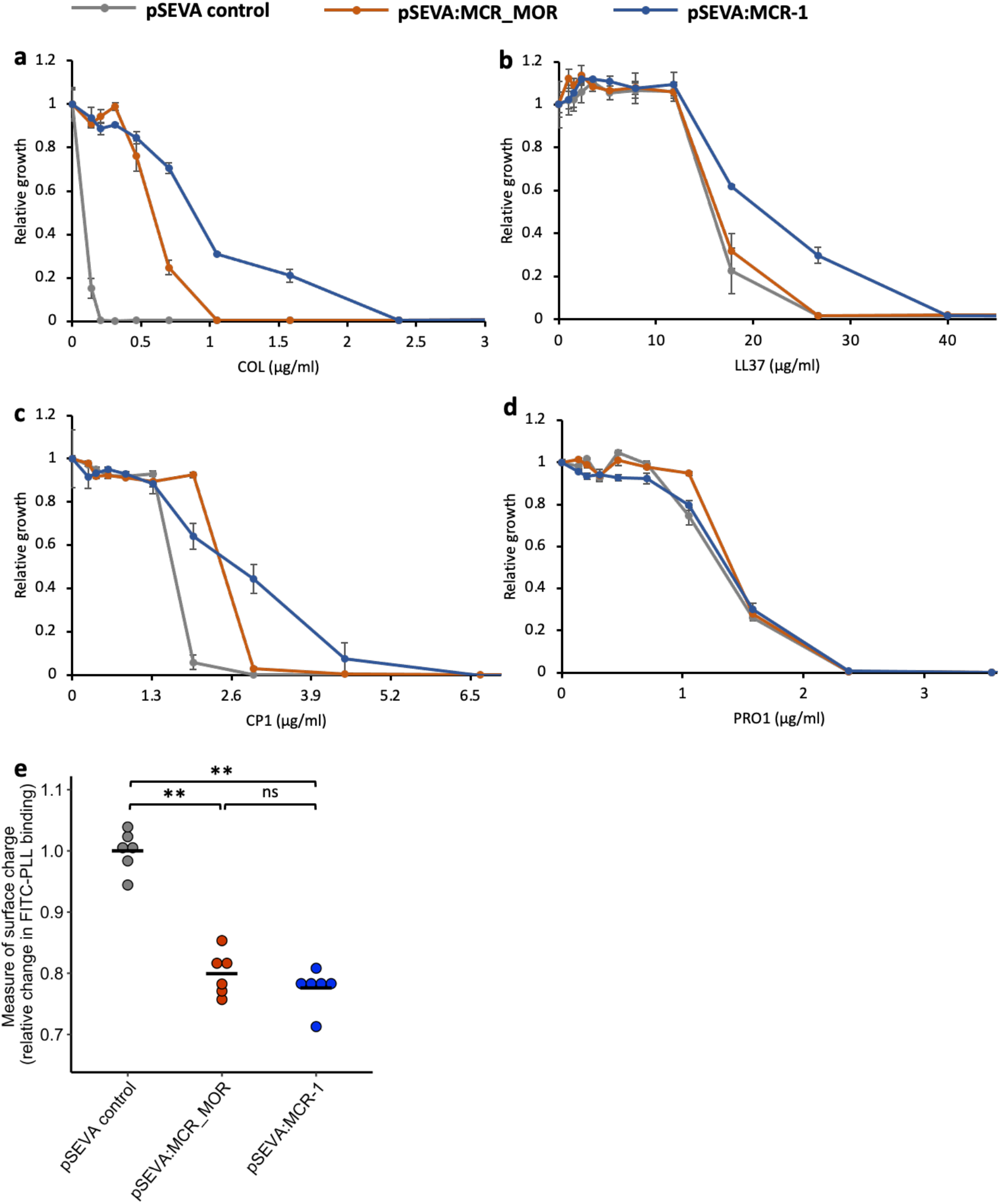
Effect of Moraxella MCR (MCR-MOR) on bacterial susceptibility to AMPs (a-d) and on cell surface charge (e). **(a-d)** AMP susceptibility of *E. coli* carrying pSEVA:MCR-1 and *Moraxella* version of MCR (pSEVA:MCR-MOR). Growth was measured as OD of AMP-treated cultures relative to AMP-free controls. Error bars indicate standard errors based on three biological replicates. **(e)** Relative cell surface charge of *E. coli* strains expressing MCR-1 and MCR-MOR compared to an empty vector control. Cell surface was determined by FITC-PLL binding assay (*n* = 6 biological replicates/strain). Statistical significance was determined by pairwise comparisons using the two-sided Mann–Whitney U tests, and double asterisks show differences with a *P* value <0.01.

Loss of negative membrane charge has been argued to play an important role in the colistin resistance provided by MCR-1. The *mcr-1* encodes for phosphoethanolamine (pEtN) transferase enzyme that facilitates the addition of phosphoethanolamine (pEtN) to the lipid A component of lipopolysaccharide (LPS), resulting in reduced binding of colistin. However, MCR-1 and MCR-MOR have similar effects on cell surface charge (Figure 3e, *P* = 0.470, two-sided Mann–Whitney U test), supporting the idea that MCR-1-mediated colistin resistance is also attributable to other factors, such as the increased protection of the cytoplasmic membrane from colisitin^46^. Given that MCR-MOR does not confer broad resistance to host AMPs, our results suggest that MCR-1 was able to evolve to increase resistance to both colistin and relevant host AMPs.

### MCR confers serum resistance and increases virulence

The above experiments focused on measuring the impact of MCR-1 on bacterial resistance to individual host AMPs. To better understand the protective role of MCR-1 in a complex host environment, we measured bacterial susceptibility to human serum, which contains a complex mixture of antimicrobials. For this assay, we selected IncI2 and IncX4 plasmids as they are the most dominant MCR-1 plasmid types^31,32^. Interestingly, these MCR plasmids conferred high levels of resistance to human serum, showing that MCR-1 is effective at providing protection against even complex mixtures of antimicrobials (Figure 4a). To rule out that the observed serum resistance is due to MCR-1 and not because of pleiotropic effects of other genes present on the plasmid, we tested serum susceptibility of a strain carrying MCR-1 knockout IncX4 plasmid. We found no difference in serum resistance between wild-type (carrying no plasmid) and strain with MCR-1 knockout plasmid, suggesting that indeed the observed serum resistance phenotype was due to MCR-1 (Figure 4b).

**Figure 4.**
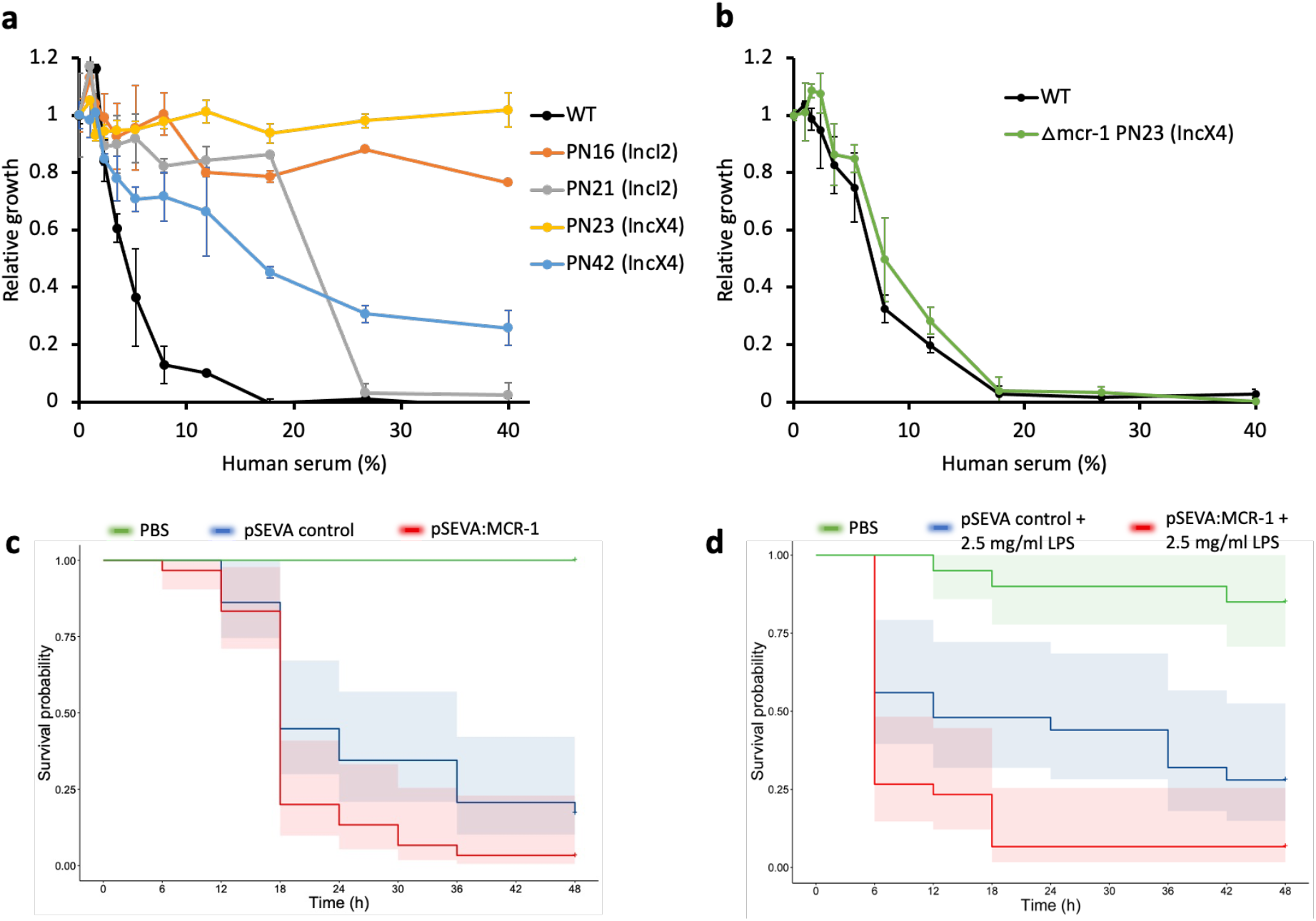
MCR confers resistance to human serum and increases bacterial virulence. **(a and b)** Bacterial susceptibility to human serum was determined by measuring bacterial growth in a medium containing human serum relative to serum-free controls (+/- s.e.m; *n* = 3 biological replicates/strain). **(a)** Serum susceptibility of the wild-type (WT) parental strain and transconjugants carrying natural MCR plasmids. **(b)** Serum susceptibility of the WT parental strain and transconjugant carrying a plasmid with a deletion of *mcr-1*. **(c and d)** Survival of *G. mellonella* larvae following injection with 5×10^7^ *E. coli* carrying pSEVA:MCR-1 or an empty vector control compared to mock-treated larvae that were injected with PBS. In **(d)** larvae were pre-treated with LPS for 24 hours prior to bacterial infection. Each experiment was performed in triplicate with 10 animals per treatment per replicate and shaded areas show 95% confidence intervals in survival probability.

These results raised the intriguing possibility that increased AMP resistance provided by MCR-1 could increase bacterial virulence by compromising host innate immunity. This is plausible as AMP resistance in pathogens has been shown to be an important virulence factor^16^. In contrast to this expectation, previous work has shown that MCR-1 plasmids actually decrease virulence in a *G. mellonella* model^34^. However, this study also showed that plasmids with identical *mcr-1* genes had differential effects on virulence, suggesting that these plasmids had effects on virulence that were unrelated to MCR-1. To directly test the impact of MCR-1 on virulence, we measured the impact of the pSEVA:MCR-1 on virulence in the *G*.*mellonella* infection model. The key advantage of this system is that pSEVA makes it possible to measure the impact of realistic levels of MCR-1 expression, while controlling for any background plasmid effects using an empty vector control. Crucially, MCR-1 carrying strain showed significantly increased killing of larvae as compared to the control strain with an empty vector, in spite of the costs associated with MCR-1 expression (Figure 4c, log-rank test, *P* = 0.024, Supplementary Figure 1).

MCR-1-mediated modification of LPS can result in reduced stimulation of macrophages and limited release of inflammatory molecules, suggesting that MCR-1 could increase virulence by reducing host immunostimulation^34^. If this is the case, then stimulating host immunity should attenuate the effect of MCR-1 on virulence. To test this idea, we measured the impact of MCR-1 on virulence in *G. mellonella* larvae that had been pre-treated with LPS, stimulating innate immunity^47^. However, MCR-1 continued to increase virulence in LPS-treated larvae, suggesting that reduced host immunostimulation was not responsible for the increased virulence associated with MCR-1 expression (Figure 4d, log-rank test *P* = 0.0074).

## Discussion

AMPs have been advocated as a potential therapeutic solution to the AMR crisis, and colistin resistance provides a unique opportunity to study the evolutionary consequences of large-scale anthropogenic AMP use. Our study shows that MCR increases bacterial fitness and resistance in the presence of AMPs from humans and agricultural animals that act as important sources of MCR carrying *E. coli* (Figure 1 and Figure 2). MCR-1 also increases resistance to human serum and virulence in an insect infection model, highlighting the threat of infections caused by MCR-*E. coli*^48^. These findings suggest that MCR-1 provides effective resistance against AMP cocktails that are found in host tissues, but it is important to emphasize that MCR mediated protection against other antimicrobials, such as lysozyme^49^, may also contribute to this protective phenotype.

One of the most important insights from this study is that anthropogenic use of AMPs (e.g. colistin) can inadvertently drive the evolution of resistance to key components of innate immunity^10,50^. Numerous AMPs are currently in clinical trials, including AMPs of human origin^4,8^, and our results highlight the importance of assessing the impact of evolved resistance to therapeutic AMP resistance on host innate immunity and bacterial virulence during pre-clinical development using sensitive and quantitative assays. It is possible, of course, that resistance to therapeutic AMPs will not be always associated with cross-resistance to host AMPs, as we observed for PROPH and PR39 (Figure 2b). However, we argue that cross-resistance to host AMPs is likely to be widespread, given that AMPs tend to share broad cellular targets and physicochemical properties (Supplementary Data 1).

MCR-1 initially spread in agricultural settings in China, where colistin was heavily used as a growth promoter. The Chinese government banned the use of colistin as a growth promoter in 2016, and this was followed by a decline in the prevalence of MCR in human, agricultural and environmental samples at a national level^51,52^. Fitness costs associated with MCR-1^34^ are likely to have played an important role in the decline of colistin resistance, but it is clear that AMPs from humans and agricultural animals provide a selective advantage for MCR-1. The doses of AMPs required to generate selection for MCR-1 resistance (∼1/2 MIC) are high compared to those that are required to select for antibiotic resistance (typically <1/10 MIC). However, AMPs achieve high concentrations in host tissues with acute or chronic inflammation^8,53^ and our results suggest that the selective benefits of AMP resistance may help to maintain MCR-1 in humans and animals, even if colistin usage remains low.

Our approach to understanding the consequences of AMP resistance evolution focused on testing the importance of the diversity of plasmid replicons that carry MCR-1. The limitation of this approach is that we tested all of these plasmids types in a single wild-type host strain. This is a limitation because the extensive genetic diversity of *E. coli* ensures that a single strain can not be assumed to be the representative of this species. An interesting avenue for further work will be to test for epistatic effects of MCR-1 across host strains. A further limitation of our study is the challenge of linking AMP resistance and virulence. Whilst the increased virulence associated with MCR carriage is consistent with the idea that MCR compromises innate immunity, it is important to emphasize that virulence is complex and multifactorial, and it is possible that increased virulence stems from changes to host tissue invasion and growth stemming from cell membrane alterations mediated by MCR-1, and not increased resistance to host immunity.

## Acknowledgements

This work was supported by grants from the Wellcome Trust (106918/Z/15Z, C.M.), the Medical Research Council (MR/S013768/1, T.W. and C.M.), the National Natural Science Foundation of China (81861138051, Y.W.), the European Research Council H2020-ERC-2014-CoG 648364–Resistance Evolution (C.P.), the European Research Council H2020-ERC-2019-PoC, 862077 Aware and Hungarian Academy of Sciences Momentum ’Célzott Lendület’ Programme LP-2017–10/2017 (C.P.), National Research, Development and Innovation Office ‘Élvonal’ Programme KKP 126506 (C.P.), National Laboratories Program, National Laboratory of Biotechnology Grant NKFIH-871-3/2020, Gazdaságfejlesztési és Innovációs Operatív Program GINOP-2.3.2–15–2016–00014 (EVOMER, C.P.), Gazdaságfejlesztési és Innovációs Operatív Program GINOP-2.3.2–15–2016–00020 (MolMedEx TUMORDNS), and National Research, Development and Innovation Office, Hungary. L.O. was supported by the Biotechnology and Biological Sciences Research Council doctoral training partnership (BB/M011224/1). P.S. was supported by the ÚNKP-21-4-New National Excellence Program of the Ministry for Innovation and Technology from the source of the National Research, Development, and Innovation Fund. M.C. was supported by the Szeged Scientists Academy under the sponsorship of the Hungarian Ministry of Innovation and Technology (FEIF/433-4/2020-ITM_SZERZ). We thank Liam P. Shaw (MacLean lab, Oxford) for helpful comments. We also thank Mei Li (Institute of Infection and Immunity, Cardiff University, UK) for assistance with the shipment of MCR-3 plasmids.

## Author contributions

P.K.J. and C.M. conceived the project and planned experiments. L.O. constructed pSEVA mutants. T.R.W., Y.Y., Q.W. and Y.W. provided MCR plasmids. P.K.J., L.O., P.S., M.C. and E.J.S. contributed to the experiments and data analysis. P.S. and M.C. carried out *Galleria* virulence assay with input from C.P. and P.K.J. P.K.J. and C.M. wrote the manuscript with input from all co-authors.

## Competing interests

The authors declare no competing interests.

## Data availability

All data generated or analysed during this study are included in this article and its Supplementary Information.

## Methods

### Bacterial strains, MCR plasmids and growth medium

All the experiments were done in *E. coli* strain J53 genetic background. All bacterial strains and MCR plasmids used in this study are listed in Supplementary Table 1 and Supplementary Table 2. Experiments were conducted in Mueller-Hinton (MH) medium and Luria-Bertani (LB) medium. All components were purchased from Sigma-Aldrich.

### AMPs

AMPs were custom synthesized by BioServ UK Ltd, except for HBD-3 and colistin. HBD-3 was custom synthesized by PeptideSynthetics UK, and colistin was purchased from Sigma-Aldrich. AMP solutions were prepared in sterile water and stored at −80°C until further use.

### Oligonucleotides

A full list of DNA oligonucleotides used in this work is provided in Supplementary Data 4. All oligos were ordered with standard desalting from Thermo Scientific.

### pSEVA:MCR-1 vector construction

A synthetic MCR-1 plasmid was constructed by cloning *mcr-1* gene into pSEVA121 plasmid^54^. The *mcr-1* gene along with its natural promoter was PCR-amplified from the natural PN16 (IncI2) plasmid using Q5® High-Fidelity DNA Polymerase (New England BioLabs). The amplified and purified *mcr-1* fragment was cloned into PCR-amplified pSEVA121 backbone using NEBuilder® HiFi DNA Assembly Master Mix according to the manufacturer’s instructions. Assembled products were then transformed into *E. coli* J53 strain using the standard electroporation method. Briefly, pSEVA121:MCR-1 plasmid carrying cells were grown overnight in MHB medium supplemented with 100 μg/ml ampicillin. Plasmid DNA isolation was performed using GeneJET Plasmid Miniprep Kit (Thermo Scientific) according to the manufacturer’s instructions. 1 μl of the purified plasmid DNA was transformed by electroporation into 50 μl of electrocompetent *E. coli* J53 cells. Electroporation was carried out with a standard protocol for a 1 mm electroporation cuvette. Cells were recovered in 1 ml SOC medium, followed by 1 h incubation at 37 °C. Different dilutions of transformant mixture were made and were plated onto petri dishes containing LB agar supplemented with 100 μg/ml ampicillin. The culture plates were incubated at 37°C for overnight.

PCR and sequence verification by Sanger sequencing were performed to ensure the presence of the correctly assembled recombinant plasmid. A full list of the primers used is given in Supplementary Data 4.

### Construction of Δ*mcr-1* PN23 (IncX4) plasmid

Gibson assembly was used to construct Δ*mcr-1* PN23 (IncX4) mutant where *mcr-1* gene was replaced by ampicillin resistance marker. The primers used for the Gibson assembly are listed in Supplementary Data 4. The overlap between fragments to be assembled was in the range of 20 to 40 bp. To avoid any mutation incorporation in the assembly, Q5® High-Fidelity 2X Master Mix (New England BioLabs) was used for PCR amplification. Five PCR fragments (leaving MCR-1 out) were generated using natural PN23 IncX4 plasmid as template DNA in Q5® High-Fidelity 2X Master Mix with corresponding primer sets (Supplementary Data 4). An ampicillin resistance marker was amplified separately.

To remove any plasmid DNA template contamination, the amplified PCR products were digested with DpnI (New England BioLabs) for 1 h at 37°C, followed by 20 min heat inactivation at 80°C. The digested PCR products were subjected to gel purification using GeneJET Gel Extraction and DNA Cleanup Micro Kit (Thermo Scientific). The gel-purified PCR products were assembled together with the ampicillin marker fragment using NEBuilder® HiFi DNA Assembly Master Mix according to the manufacturer’s instructions. The resulting assembled plasmid DNA was transformed into *E. coli* strain MG1655, rather transforming directly into *E. coli* J53. This extra step was to ensure efficient transformation of the assembled plasmid. *E. coli* MG1655 is well lab-adapted strain and shows high transformation efficiency, especially for large plasmids. The transformants were selected on LB agar containing ampicillin 100 μg/ml. The presence and right orientation of all six fragments were confirmed by PCR amplification of fragments junction. Similarly, the absence of *mcr-1* gene was also confirmed by PCR. Following the confirmation of the Δ*mcr-1* PN23 (IncX4) plasmid, a conjugation experiment was carried out to transfer Δ*mcr-1* PN23 (IncX4) plasmid into *E. coli* J53.

### Conjugation experiments

Conjugation experiments were carried out in LB broth medium at 37°C using *E. coli* strain J53 as the recipient and MCR-1-positive *E. coli* (MCRPEC) natural strains as the donor. The overnight grown cultures of both the donor and recipient strain were washed with fresh LB medium and mixed at a 1:1 ratio. The mixed culture was incubated at 37°C for overnight without shaking. Transconjugants were selected on LB agar containing 150 μg/ml sodium azide and 2 μg/ml colistin. In the case of mcr-knockout plasmid mutant (Δ*mcr-1* PN23 IncX4), *E. coli* MG1655 was used as the donor and the transconjugants were selected on 150 μg/ml sodium azide and 100 μg/ml ampicillin. The presence of plasmids in transconjugants was confirmed by PCR.

### Construction of pSEVA:MCR-MOR plasmid

*Moraxella* species have been identified as potential sources of MCR-1^32,43^. To study the Moraxella version of MCR (MCR-MOR), we custom synthesized (Twist Bioscience) MCR-MOR gene (*Moraxella osloensis*, GenBank: AXE82_07515) and cloned this gene into pSEVA121 plasmid using Gibson assembly method. For cloning, the MCR-MOR fragment (insert DNA 1709 bp) and pSEVA backbone (vector DNA 4001 bp) containing ampicillin resistance marker were amplified by PCR with corresponding primers (Supplementary Data 4) in Q5® High-Fidelity 2X Master Mix (New England BioLabs). Both the insert (MCR-MOR) and vector fragments were gel purified using GeneJET Gel Extraction and DNA Cleanup Micro Kit (Thermo Scientific). The gel-purified PCR products were assembled together using NEBuilder® HiFi DNA Assembly Master Mix (New England BioLabs) according to the manufacturer’s instructions. Following the assembly, 2ul of the assembly mixture was transformed into *E. coli* strain J53 and transformants were selected on LB agar containing 100 μg/ml ampicillin. The assembly of pSEVA MCR-MOR plasmid was verified by PCR.

### Physicochemical properties of AMPs

Protein amino acid frequencies and the fraction of polar and non-polar amino acids were counted with an in-house R script. PepCalc (Innovagen) calculator was used to calculate the net charge. Isoelectric point and hydrophobicity were calculated using Peptide Analyzing Tool (Thermo Scientific). Percentage of the disordered region, beta-strand region, coiled structure and alpha-helical region was calculated with Pasta 2.0. The ExPasy ProtParam tool was used for calculating aliphatic index and hydropathicity. Aggregation hotspots were calculated by AggreScan.

### Determination of minimum inhibitory concentration (MIC)

Minimum inhibitory concentrations (MICs) were determined with a standard serial broth dilution technique with a minor modification that we previously optimized for AMPs^40^. Specifically, smaller AMP concentration steps were used (typically 1.2-1.5-fold) because AMPs have steeper dose-response curves than standard antibiotics^3,4^, and therefore bigger concentration steps (such as two-fold dilutions) can not capture 90% growth inhibitions (i.e., MIC). 10-steps serial dilution was prepared in fresh MHB medium in 96-well microtiter plates where AMP was represented in 9 different concentrations. Three wells contained only medium to monitor the growth in the absence of AMP. Bacterial strains were grown in MHB medium supplemented with appropriate antibiotic (100 μg/ml ampicillin for *E. coli* pSEVA MCR-1 and 1 μg/ml colistin for MCR natural plasmid) at 30°C for overnight. Following overnight incubation, approximately 5×10^5^ cells were inoculated into the wells of the 96-well microtiter plate. We used three independent replicates for each strain and the corresponding control. The top and bottom row in the 96-well plate were filled with MHB medium to obtain the background OD value of the medium. Plates were incubated at 30 °C with continuous shaking at 250 rpm. After 20–24 h of incubation, OD_600_ values were measured in a microplate reader (Biotek Synergy 2). After background subtraction, MIC was defined as the lowest concentration of AMP where the OD_600_ <0.05. Bacterial susceptibility to human serum was also measured using the similar MIC assay described above. Human serum was purchased from Sigma.

### Membrane surface charge measurement

To measure bacterial membrane surface charge, we carried out a fluorescein isothiocyanate-labelled poly-L-lysine (FITC-PLL) (Sigma) binding assay. FITC-PLL is a polycationic molecule that binds to an anionic lipid membrane in a charge-dependent manner and is used to investigate the interaction between cationic peptides and charged lipid bilayer membranes^55^. The assay was performed as previously described^5,40^. Briefly, bacterial cells were grown overnight in MHB medium, centrifuged and washed twice with 1X PBS buffer (pH 7.4). The washed bacterial cells were re-suspended in 1× PBS buffer to a final OD_600_ of 0.1. A freshly prepared FITC-PLL solution was added to the bacterial suspension at a final concentration of 6.5 μg/ml. The suspension was incubated at room temperature for 10 min, and pelleted by centrifugation. The remaining amount of FITC-PLL in the supernatant was determined fluorometrically (excitation at 500 nm and emission at 530 nm) with or without bacterial exposure. The quantity of bound molecules was calculated from the difference between these values. A lower binding of FITC-PLL indicates a less net negative surface charge of the outer bacterial membrane.

### *In vitro* competition assay

To directly test the selective fitness benefits of MCR-1, we carried out *in vitro* competition experiment using a flow cytometry-based sensitive and reproducible method developed in our lab^34,56,57^. Flow cytometry was performed on an Accuri C6 (Becton Dickenson, Biosciences, UK). We measured the competitive fitness of *E. coli* strain J53 harbouring pSEVA MCR-1 in the absence and presence of an AMP. For this assay, we randomly selected five AMPs and colistin. *E. coli* harbouring pSEVA plasmid without MCR-1 (called pSEVA <u>e</u>mpty <u>v</u>ector (EV)) was used as a control to calculate the relative fitness of *E. coli* pSEVA:MCR-1. These strains were competed against a GFP-labelled *E. coli* strain J53 to measure the relative fitness (see Supplementary Figure 4). All competitions were carried out in MHB medium with six biological replicates per strain, as previously described^34,56^. Briefly, the bacterial cells were grown in MHB medium supplemented with 100 ug/ml ampicillin at 30°C for overnight. The overnight grown cultures were washed with filtered PBS buffer to remove any antibiotic residues. The washed cells were diluted into a fresh MHB medium and mixed approximately at 1:1 ratio with GFP-labelled *E. coli* J53. Before starting the competition, the total cell density in the competition mix was around half million cells, as we also used for MIC assay. The initial density of fluorescent and non-fluorescent cells was estimated in the mix using medium flow rate, recoding 10,000 events and discarding events with forward scatter (FSC) <10,000 and side scatter (SSC) <8000. After confirming the actual ratio close to 1:1, the competition plates were incubated at 30°C with shaking at 250rpm. After overnight incubation, the competition mix was diluted in PBS buffer and cell densities were adjusted around 1000 per microlitre. The final density of fluorescent and non-fluorescent cells was estimated in the competition mix. Using the initial and final density of fluorescent and non-fluorescent cells, the relative fitness was calculated as described below:

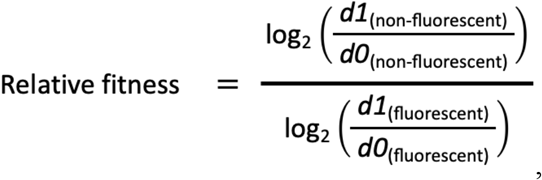

Where *d0* and *d1* represent cell density before and after the competition, respectively. Using this formula, the fitness of *E. coli* pSEVA:MCR-1 and *E. coli* pSEVA EV control was calculated (relative to GFP-labelled strain). In Figure 1, we expressed the fitness of *E. coli* pSEVA:MCR-1 strain relative to the control strain (i.e., *E. coli* pSEVA EV), and followed the procedure of error propagation to account for the uncertainty of the estimates:

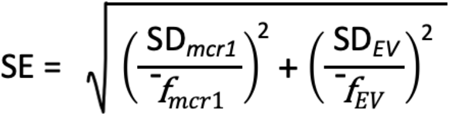

where ^-^*f* and SD are a mean estimate and its standard deviation for each corresponding strain based on six biological replicates. MCR1 and EV represent *E. coli* J53 carrying pSEVA:MCR-1 and *E. coli* J53 carrying empty vector control strain, respectively.

### *In vivo* virulence assay

Age and weight defined TruLarv™ *Galleria mellonella* caterpillars were obtained in bulk from BioSystems Technology (Exeter, United Kingdom) and stored at 15°C in absence of food. *E. coli* J53 pSEVA:MCR-1 and empty vector control strain was grown overnight in MHB broth and washed twice with sterile PBS. In the case of every experiment, treatment solutions were injected into the hemocoels of the larvae via the first right proleg using 10 μl Hamilton syringes (Reno, Nevada, U.S.A.). Larvae were incubated in petri dishes lined with filter paper at 37°C for 48 h and survival was documented every 6 hours. Insects were considered dead if they failed to respond to touch. Pretreatment was administered approximately 24 hours before bacterial injection, and in this time period the survival of the animals was not recorded. Before bacterial injection, the dead or sick animals were excluded from further experiments.

In order to establish the inoculum required to kill *G. mellonella* over 48 hours, 10 caterpillars were inoculated with 10 μl of bacterial suspensions containing 10^4^, 10^5^, 10^6^, 10^7^ and 10^8^ CFU/larva of *E. coli* strain carrying pSEVA empty vector control in PBS (data not shown). CFU number was verified by viable bacterial counts on MHB agar. Based on this preliminary experiment 5*10^7^ and 1*10^8^ were determined as the ideal inoculum sizes to kill *G. mellonella* larvae.

LPS from pathogenic bacterial strain *Escherichia coli* O111:B4 was purchased from Sigma Aldrich (Merck KGaA, Darmstadt, Germany) that has been shown to stimulate host innate immunity^47^. LPS solutions from powder were prepared fresh by dissolving the powder in 1X PBS and the solution was sterilized by heating at 80°C for at least 30 minutes. LPS pretreatment was administered similarly to bacterial treatment into the left first proleg approximately 24 hours before bacterial injection. In this time period the survival of the animals was not continuously recorded. Before bacterial injection, the dead or sick animals were excluded from further experiments. In order to establish an ideal treatment dose of LPS, a dose-response experiment was performed with 1.25, 2.5, 5, 10 and 20 mg/ml LPS solution used for pretreatment (data not shown). Larvae were injected with 10 μl of each dose of LPS independently. In the case of animals injected with only LPS in the absence of bacteria, the survival of the animals was not affected, proving that LPS in itself has no significant toxic effects at the tested concentrations. In the case of injecting the animals with bacteria, LPS caused a very severe reaction and swift animal death. Because of that, the relatively small treatment dose of 2.5 mg/ml was chosen for the final experiment.

All experimental data were visualized with Kaplan-Meier survival curves, utilizing R packages *survival, survminer* and *ggsurvplot*. P-values in comparison of treatment groups within experiments were generated by these packages utilizing a standard Log-Rank test. P-values comparing results between experiments were obtained by comparing hazard ratios between the treatment lines based on the Cox proportional-hazards model.

## Supplementary information

### Supplementary Figures

**Supplementary Figure 1.**
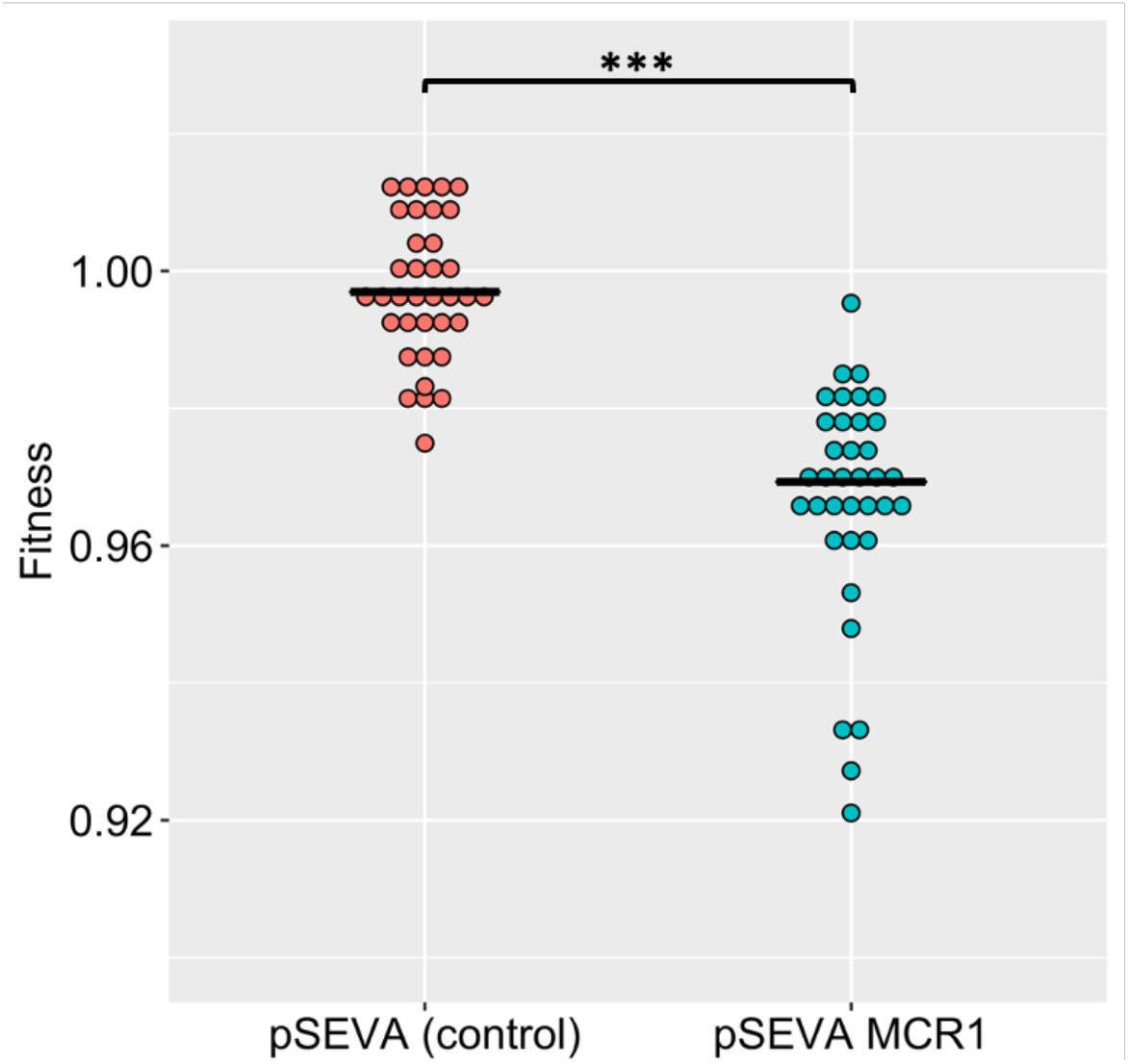
MCR-1 imposes a significant fitness burden in the absence of an AMP. (*P* = 1.174e-15, from two-sided Mann–Whitney U test, *n* = 36 for each genotype). This figure shows the combined fitness data for colistin and five host AMPs used for *in vitro* competition assay (see Figure 1). Fitness was calculated by competing the pSEVA plasmid strains against GFP-labelled *E. coli* (see Methods).

**Supplementary Figure 2.**
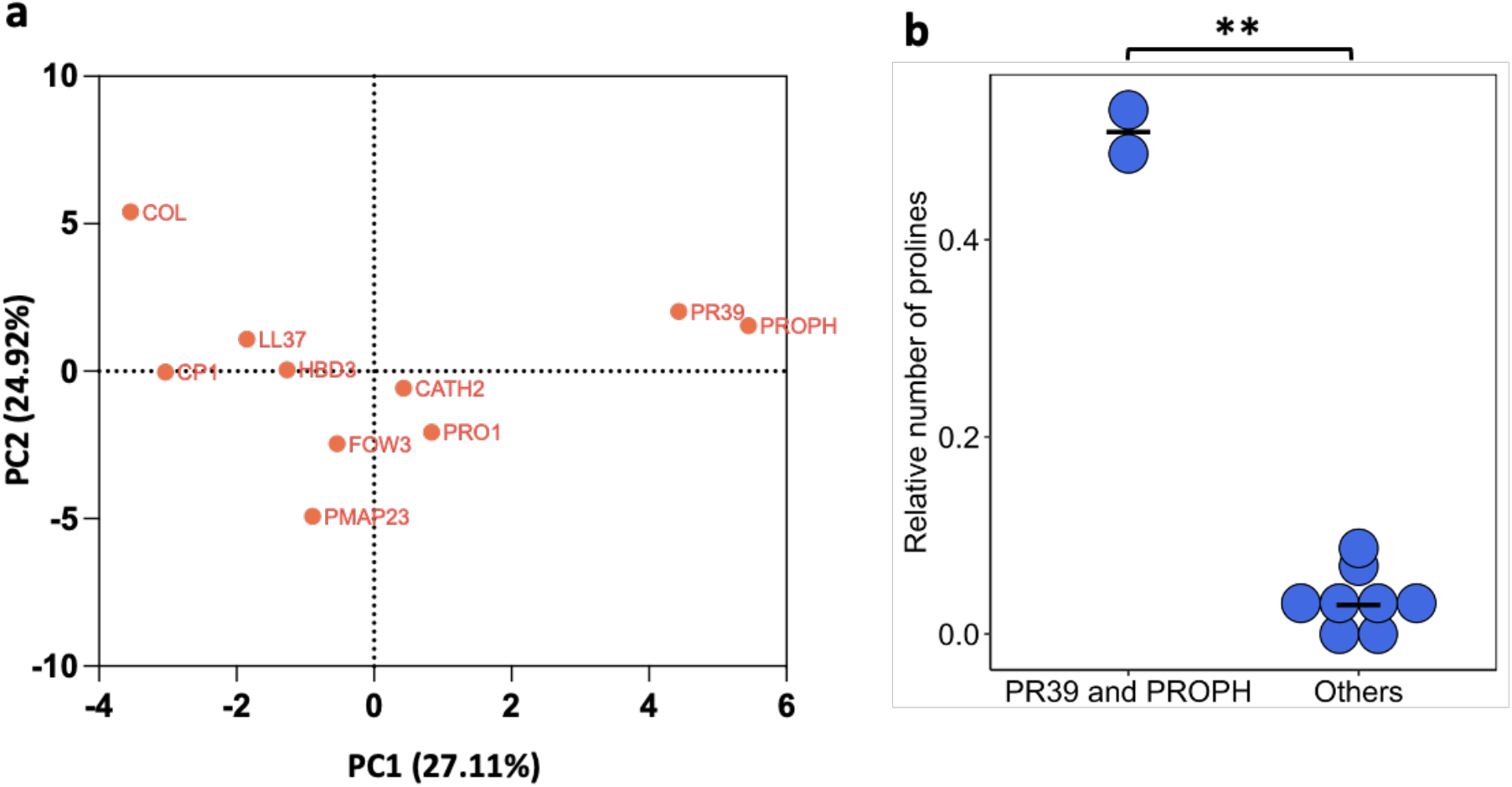
PR39 and PROPH differ from other AMPs in their physicochemical properties. **(a)** Principle component analysis (PCA) of the physicochemical properties of AMPs. **(b)** PR39 and PROPH peptides can be distinguished from other AMPs based on their higher proline content (significant differences from Welch Two Sample t-test, *P* = 0.0086, *n* = 2 and *n* = 8 for PR39 and PROPH, and others, respectively).

**Supplementary Figure 3.**
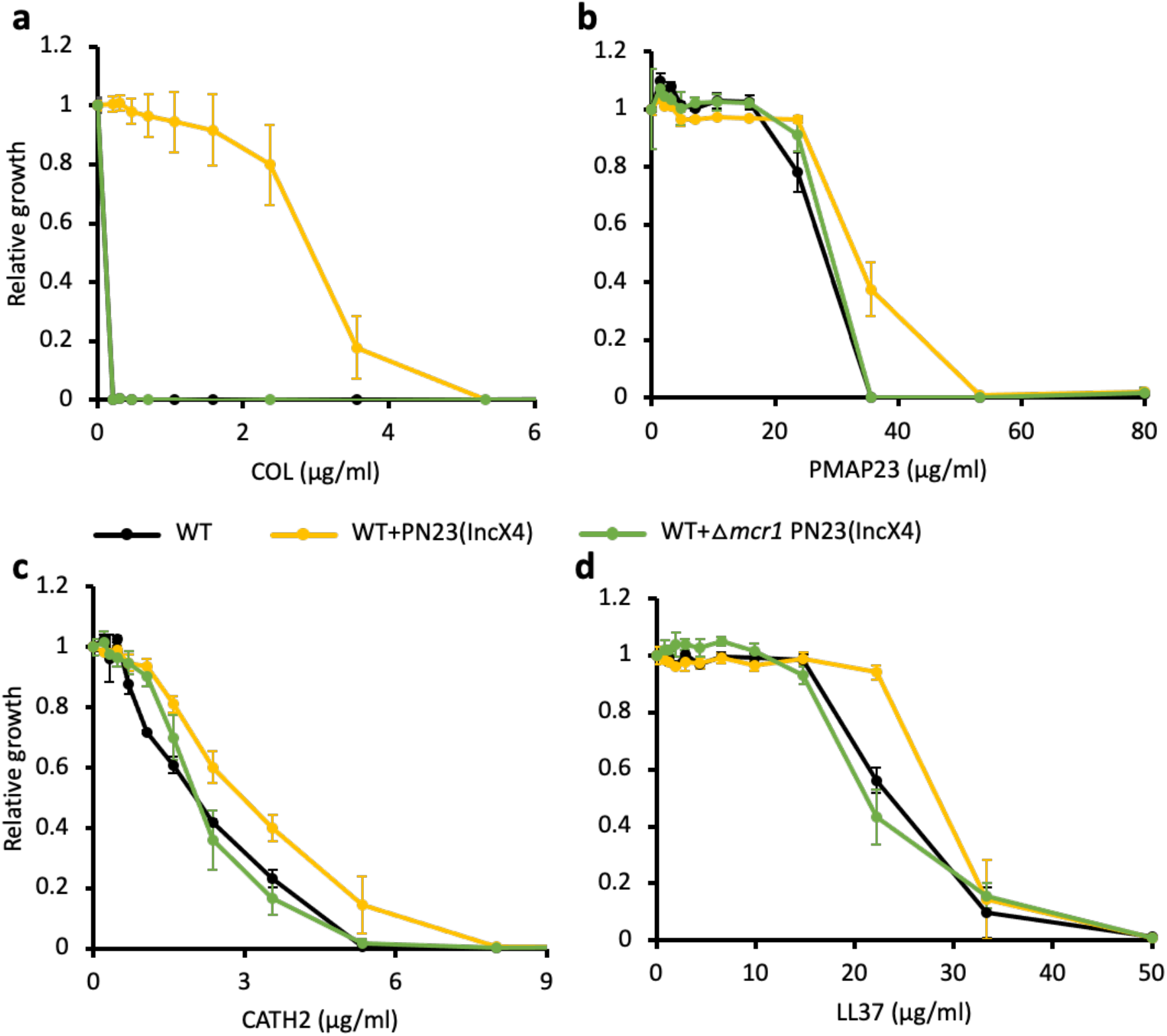
Deletion of MCR-1 from IncX4 natural plasmid results in AMP susceptibility similar to the wild-type (WT) control strain. Growth is shown relative to growth in the absence of the given AMP (y-axis). Error bars indicate standard errors based on three biological replicates. Black line – WT control strain; Yellow line-WT strain carrying IncX4 plasmid with *mcr-1* gene; Green line – WT strain carrying IncX4 plasmid without *mcr-1* gene.

**Supplementary Figure 4.**
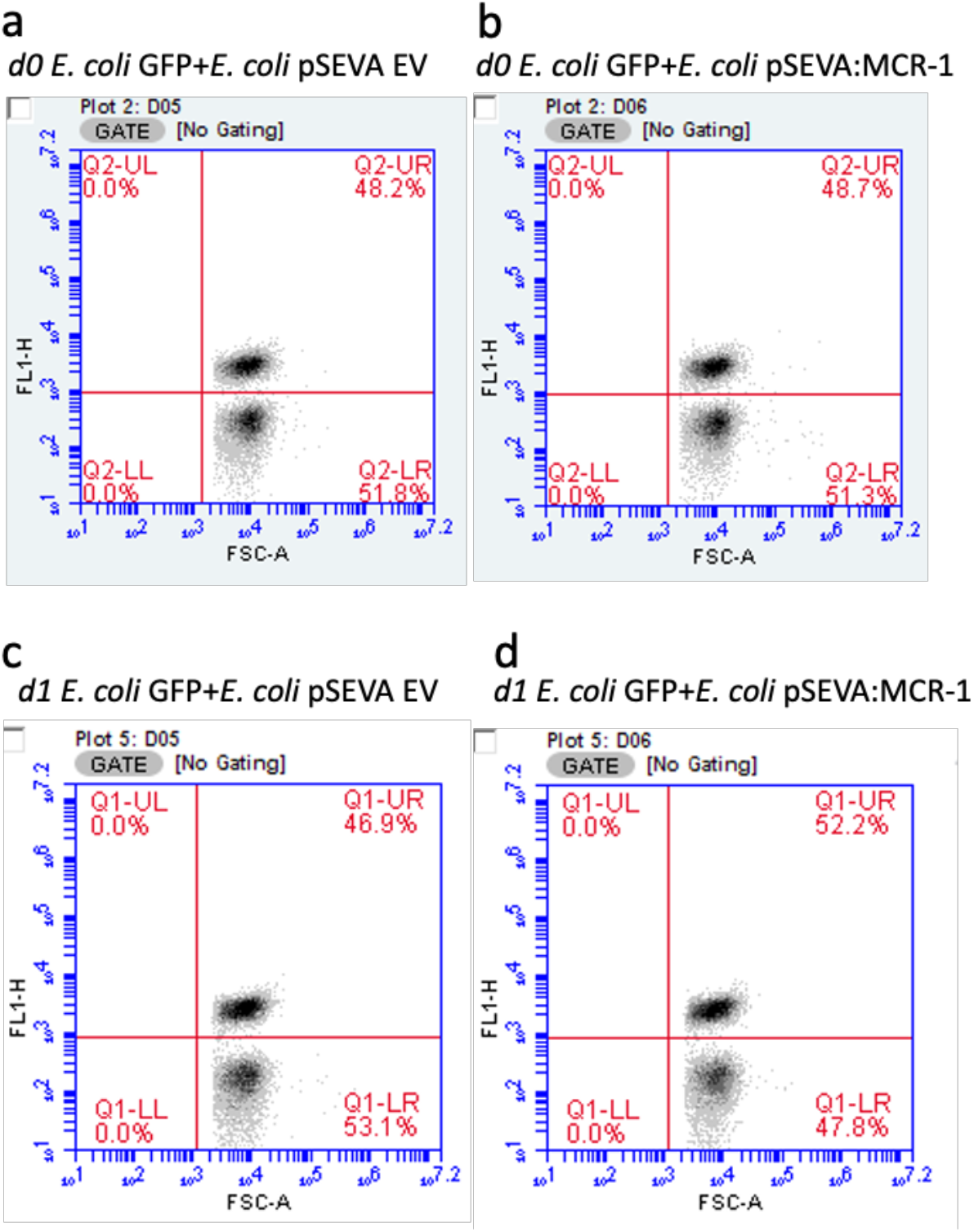
Gating strategy to analyse the selective fitness benefits of MCR-1. Figures show cell density of fluorescent (GFP-labelled *E. coli;* gate - UR) and non-fluorescent cells (*E. coli* pSEVA EV or *E. coli* pSEVA:MCR-1; gate - LR) before (a and b) and after the competition (c and d) in the absence of an AMP. Cells were gated using FL1-H and forward scatter properties. Data are provided in Supplementary Data 2.

### Supplementary Tables

**Supplementary Table 1.**
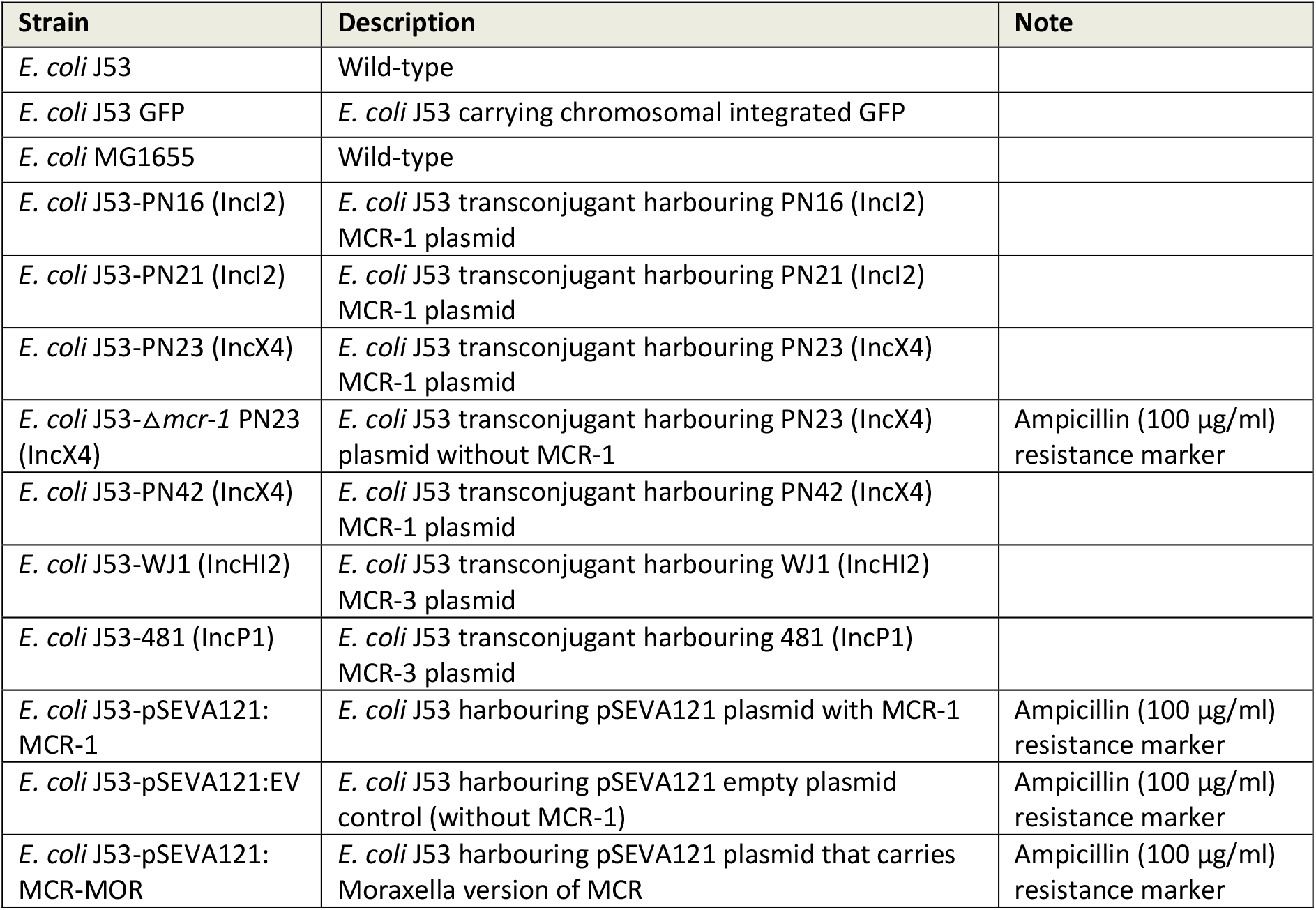
List of bacterial strains used in this study.

| Strain | Description | Note |
| --- | --- | --- |
| *E. coli* J53 | Wild-type |  |
| *E. coli* J53 GFP | *E. coli* J53 carrying chromosomal integrated GFP |  |
| *E. coli* MG1655 | Wild-type |  |
| *E. coli* J53-PN16 (IncI2) | *E. coli* J53 transconjugant harbouring PN16 (IncI2) MCR-1 plasmid |  |
| *E. coli* J53-PN21 (IncI2) | *E. coli* J53 transconjugant harbouring PN21 (IncI2) MCR-1 plasmid |  |
| *E. coli* J53-PN23 (IncX4) | *E. coli* J53 transconjugant harbouring PN23 (IncX4) MCR-1 plasmid |  |
| *E. coli* J53-$\Delta mcr-1$ PN23 (IncX4) | *E. coli* J53 transconjugant harbouring PN23 (IncX4) plasmid without MCR-1 | Ampicillin (100 µg/ml) resistance marker |
| *E. coli* J53-PN42 (IncX4) | *E. coli* J53 transconjugant harbouring PN42 (IncX4) MCR-1 plasmid |  |
| *E. coli* J53-WJ1 (IncHI2) | *E. coli* J53 transconjugant harbouring WJ1 (IncHI2) MCR-3 plasmid |  |
| *E. coli* J53-481 (IncP1) | *E. coli* J53 transconjugant harbouring 481 (IncP1) MCR-3 plasmid |  |
| *E. coli* J53-pSEVA121: MCR-1 | *E. coli* J53 harbouring pSEVA121 plasmid with MCR-1 | Ampicillin (100 µg/ml) resistance marker |
| *E. coli* J53-pSEVA121:EV | *E. coli* J53 harbouring pSEVA121 empty plasmid control (without MCR-1) | Ampicillin (100 µg/ml) resistance marker |
| *E. coli* J53-pSEVA121: MCR-MOR | *E. coli* J53 harbouring pSEVA121 plasmid that carries Moraxella version of MCR | Ampicillin (100 µg/ml) resistance marker |

**Supplementary Table 2.** Plasmids used in this study.

| Plasmid | *mcr* gene | Size (bp) | Marker gene | Note | Source |
| --- | --- | --- | --- | --- | --- |
| PN16 (IncI2) | *mcr-1* | 60488 | - | MCR-1 natural plasmid belonging to IncI2 incompatibility group | Yang et al. Nat Commun 2017 |
| PN21 (IncI2) | *mcr-1* | 60 989 | - | MCR-1 natural plasmid belonging to IncI2 incompatibility group | Yang et al. Nat Commun 2017 |
| PN23 (IncX4) | *mcr-1* | 33858 | - | MCR-1 natural plasmid belonging to IncX4 incompatibility group | Yang et al. Nat Commun 2017 |
| Δ*mcr-1* PN23 (IncX4) | - | 33221 | Ampicillin (100µg/ml) | MCR-1 deletion mutant of PN23 (IncX4) | This study |
| PN42 (IncX4) | *mcr-1* | 32995 | - | MCR-1 natural plasmid belonging to IncX4 incompatibility group | Yang et al. Nat Commun 2017 |
| WJ1 (IncHI2) | *mcr-3* | 266119 | - | MCR-3 natural plasmid belonging to IncHI2 incompatibility group | Yin et al. mBio 2017 |
| 481 (IncP1) | *mcr-3* | 53660 | - | MCR-3 natural plasmid belonging to IncP1 incompatibility group | Wang et al. Veterinary Microbiology 2019 |
| pSEVA121:EV | - | 3924 | Ampicillin (100µg/ml) | pSEVA121 synthetic plasmid without MCR-1 | Silva-Rocha et al. NAR 2013 |
| pSEVA121:MCR-1 | *mcr-1* | 5659 | Ampicillin (100µg/ml) | pSEVA121 synthetic plasmid carrying MCR-1 and promoter | This study |
| pSEVA121:MCR-MOR | *Moraxella* version of MCR | 5670 | Ampicillin (100µg/ml) | pSEVA121 synthetic plasmid carrying *Moraxella* version of MCR | This study |

### Supplementary Data files

**Supplementary Data 1. List of the physicochemical and mechanistic properties of the AMPs used in this study**.

Provided in a separate Excel spreadsheet.

**Supplementary Data 2. Competitive fitness data**

Provided in a separate Excel spreadsheet.

**Supplementary Data 3. Antimicrobial susceptibility of MCR-*E. coli* to AMPs**

Provided in a separate Excel spreadsheet.

**Supplementary Data 4. List of oligonucleotides used in this study**.

Provided in a separate Excel spreadsheet.

## Notes

### Competing Interest Statement

The authors have declared no competing interest.

## References

1. Zasloff, M. Antimicrobial peptides of multicellular organisms. Nature 415, 389–395 (2002).

2. Yeung, A. T. Y., Gellatly, S. L. & Hancock, R. E. W. Multifunctional cationic host defence peptides and their clinical applications. Cell. Mol. Life Sci. 68, (2011).

3. Yu, G., Baeder, D. Y., Regoes, R. R. & Rolff, J. Predicting drug resistance evolution: insights from antimicrobial peptides and antibiotics. Proc. R. Soc. B Biol. Sci. 285, 20172687 (2018).

4. Lazzaro, B. P., Zasloff, M. & Rolff, J. Antimicrobial peptides: Application informed by evolution. Science (80-.). 368, eaau5480 (2020).

5. Spohn, R. et al. Integrated evolutionary analysis reveals antimicrobial peptides with limited resistance. Nat. Commun. 10, 4538 (2019).

6. Jangir, P. K., Ogunlana, L. & MacLean, R. C. Evolutionary constraints on the acquisition of antimicrobial peptide resistance in bacterial pathogens. Trends Microbiol. (2021) doi:10.1016/j.tim.2021.03.007.

7. Hancock, R. E. W. & Sahl, H. -G. Antimicrobial and host-defense peptides as new anti-infective therapeutic strategies. Nat. Biotechnol. 24, 1551–1557 (2006).

8. Mookherjee, N., Anderson, M. A., Haagsman, H. P. & Davidson, D. J. Antimicrobial host defence peptides: functions and clinical potential. Nat. Rev. Drug Discov. 19, 311–332 (2020).

9. Magana, M. et al. The value of antimicrobial peptides in the age of resistance. Lancet Infect. Dis. 20, e216–e230 (2020).

10. Habets, M.G.J.L. & Brockhurst, M. A. Therapeutic antimicrobial peptides may compromise natural immunity. Biol. Lett. (2012) doi:10.1098/rsbl.2011.1203.

11. Kubicek-Sutherland, J. Z. et al. Antimicrobial peptide exposure selects for Staphylococcus aureus resistance to human defence peptides. J. Antimicrob. Chemother. 72, 115–127 (2017).

12. Fleitas, O. & Franco, O. L. Induced bacterial cross-resistance toward host antimicrobial peptides: A worrying phenomenon. Frontiers in Microbiology (2016) doi:10.3389/fmicb.2016.00381.

13. Andersson, D. I., Hughes, D. & Kubicek-Sutherland, J. Z. Mechanisms and consequences of bacterial resistance to antimicrobial peptides. Drug Resist. Updat. 26, 43–57 (2016).

14. Salzman, N. H. et al. Enteric defensins are essential regulators of intestinal microbial ecology. Nat. Immunol. 11, 76–83 (2010).

15. Ostaff, M. J., Stange, E. F. & Wehkamp, J. Antimicrobial peptides and gut microbiota in homeostasis and pathology. EMBO Mol. Med. 5, 1465–83 (2013).

16. Groisman, E. A., Parra-Lopez, C., Salcedo, M., Lippst, C. J. & Heffron, F. Resistance to host antimicrobial peptides is necessary for Salmonella virulence. Genetics vol. 89 (1992).

17. Kidd, T. J. et al. A Klebsiella pneumoniae antibiotic resistance mechanism that subdues host defences and promotes virulence. EMBO Mol. Med. 9, 430–447 (2017).

18. Pál, C., Papp, B. & Lázár, V. Collateral sensitivity of antibiotic-resistant microbes. Trends Microbiol. 23, 401–407 (2015).

19. Imamovic, L. & Sommer, M. O. A. Use of Collateral Sensitivity Networks to Design Drug Cycling Protocols That Avoid Resistance Development. Sci. Transl. Med. 5, 204ra132–204ra132 (2013).

20. Barbosa, C., Roemhild, R., Rosenstiel, P. & Schulenburg, H. Evolutionary stability of collateral sensitivity to antibiotics in the model pathogen pseudomonas aeruginosa. Elife 8, 1–22 (2019).

21. Partridge, S. R., Kwong, S. M., Firth, N. & Jensen, S. O. Mobile Genetic Elements Associated with Antimicrobial Resistance. Clin. Microbiol. Rev. 31, (2018).

22. MacLean, R. C. & San Millan, A. The evolution of antibiotic resistance. Science (80-.). 365, 1082–1083 (2019).

23. Rodríguez-Rojas, A., Makarova, O., Müller, U. & Rolff, J. Cationic Peptides Facilitate Iron-induced Mutagenesis in Bacteria. PLOS Genet. 11, e1005546 (2015).

24. Vaara, M. Agents that increase the permeability of the outer membrane. Microbiol. Rev. 56, 395–411 (1992).

25. Wang, Y. et al. Prevalence, risk factors, outcomes, and molecular epidemiology of mcr-1-positive Enterobacteriaceae in patients and healthy adults from China: an epidemiological and clinical study. Lancet Infect. Dis. 17, 390–399 (2017).

26. Li, J. et al. Colistin: the re-emerging antibiotic for multidrug-resistant Gram-negative bacterial infections. Lancet Infect. Dis. 6, 589–601 (2006).

27. Jochumsen, N. et al. The evolution of antimicrobial peptide resistance in Pseudomonas aeruginosa is shaped by strong epistatic interactions. Nat. Commun. 7, 13002 (2016).

28. Sun, B. et al. New Mutations Involved in Colistin Resistance in Acinetobacter baumannii. mSphere 5, (2020).

29. Snitkin, E. S. et al. Genomic insights into the fate of colistin resistance and Acinetobacter baumannii during patient treatment. Genome Res. 23, 1155–1162 (2013).

30. Liu, Y.-Y. et al. Emergence of plasmid-mediated colistin resistance mechanism MCR-1 in animals and human beings in China: a microbiological and molecular biological study. Lancet Infect. Dis. 16, 161–168 (2016).

31. Wang, R. et al. The global distribution and spread of the mobilized colistin resistance gene mcr-1. Nat. Commun. (2018) doi:10.1038/s41467-018-03205-z.

32. Sun, J., Zhang, H., Liu, Y.-H. & Feng, Y. Towards Understanding MCR-like Colistin Resistance. Trends Microbiol. 26, 794–808 (2018).

33. Gao, R. et al. Dissemination and Mechanism for the MCR-1 Colistin Resistance. PLOS Pathog. 12, e1005957 (2016).

34. Yang, Q. et al. Balancing mcr-1 expression and bacterial survival is a delicate equilibrium between essential cellular defence mechanisms. Nat. Commun. 8, 2054 (2017).

35. Lázár, V. et al. Antibiotic-resistant bacteria show widespread collateral sensitivity to antimicrobial peptides. Nat. Microbiol. 3, 718–731 (2018).

36. Barlow, P. G. et al. The Human Cathelicidin LL-37 Preferentially Promotes Apoptosis of Infected Airway Epithelium. Am. J. Respir. Cell Mol. Biol. 43, (2010).

37. Srakaew, N. et al. Antimicrobial host defence peptide, LL-37, as a potential vaginal contraceptive. Hum. Reprod. 29, (2014).

38. Zasloff, M. Antimicrobial Peptides in Health and Disease. N. Engl. J. Med. 347, (2002).

39. Zhang, L. & Gallo, R. L. Antimicrobial peptides. Curr. Biol. 26, (2016).

40. Kintses, B. et al. Chemical-genetic profiling reveals limited cross-resistance between antimicrobial peptides with different modes of action. Nat. Commun. 10, 5731 (2019).

41. Scocchi, M., Tossi, A. & Gennaro, R. Proline-rich antimicrobial peptides: converging to a non-lytic mechanism of action. Cell. Mol. Life Sci. 68, (2011).

42. Gerstel, U. et al. Hornerin contains a Linked Series of Ribosome-Targeting Peptide Antibiotics. Sci. Rep. 8, (2018).

43. Kieffer, N., Nordmann, P. & Poirel, L. Moraxella Species as Potential Sources of MCR-Like Polymyxin Resistance Determinants. Antimicrob. Agents Chemother. 61, (2017).

44. Wei, W. et al. Defining ICR-Mo, an intrinsic colistin resistance determinant from Moraxella osloensis. PLOS Genet. 14, (2018).

45. AbuOun, M. et al. mcr-1 and mcr-2 (mcr-6.1) variant genes identified in Moraxella species isolated from pigs in Great Britain from 2014 to 2015. J. Antimicrob. Chemother. 72, (2017).

46. Sabnis, A. et al. Colistin kills bacteria by targeting lipopolysaccharide in the cytoplasmic membrane. Elife 10, (2021).

47. Mukherjee, K. et al. Galleria mellonella as a model system for studying Listeria pathogenesis. Appl. Environ. Microbiol. 76, 310–317 (2010).

48. Yin, W. et al. Mobile Colistin Resistance Enzyme MCR-3 Facilitates Bacterial Evasion of Host Phagocytosis. Adv. Sci. 2101336, 1–15 (2021).

49. Sherman, E. X., Hufnagel, D. A. & Weiss, D. S. MCR -1 confers cross-resistance to lysozyme. Lancet Infect. Dis. 16, 1226–1227 (2016).

50. Gabriel G Perron, M. Z. and G. B. Experimental evolution of resistance to an antimicrobial peptide. Proc. R. Soc. B. 273, 251–256 (2005).

51. Shen, C. et al. Dynamics of mcr-1 prevalence and mcr-1-positive Escherichia coli after the cessation of colistin use as a feed additive for animals in China: a prospective cross-sectional and whole genome sequencing-based molecular epidemiological study. The Lancet Microbe 1, e34–e43 (2020).

52. Wang, Y. et al. Changes in colistin resistance and mcr-1 abundance in Escherichia coli of animal and human origins following the ban of colistin-positive additives in China: an epidemiological comparative study. Lancet Infect. Dis. 20, (2020).

53. Fahlgren, A., Hammarström, S., Danielsson, Å. & Hammarström, M.-L. Increased expression of antimicrobial peptides and lysozyme in colonic epithelial cells of patients with ulcerative colitis. Clin. Exp. Immunol. 131, (2003).

54. Silva-Rocha, R. et al. The Standard European Vector Architecture (SEVA): a coherent platform for the analysis and deployment of complex prokaryotic phenotypes. Nucleic Acids Res. 41, (2013).

55. Rossetti, F. F. et al. Interaction of Poly(L-Lysine)-g-Poly(Ethylene Glycol) with Supported Phospholipid Bilayers. Biophys. J. 87, (2004).

56. San Millan, A., Escudero, J. A., Gifford, D. R., Mazel, D. & MacLean, R. C. Multicopy plasmids potentiate the evolution of antibiotic resistance in bacteria. Nat. Ecol. Evol. 1, (2017).

57. Gifford, D. R. et al. Identifying and exploiting genes that potentiate the evolution of antibiotic resistance. Nat. Ecol. Evol. 2, (2018).

## References for supplementary information

1. Yang, Q. et al. Balancing mcr-1 expression and bacterial survival is a delicate equilibrium between essential cellular defence mechanisms. Nat. Commun. 8, 2054 (2017).

2. Yin, W. et al. Novel Plasmid-Mediated Colistin Resistance Gene mcr-3 in Escherichia coli. (2017) doi:10.1128/mBio.

3. Wang, Z. et al. Genetic environment of colistin resistance genes mcr-1 and mcr-3 in Escherichia coli from one pig farm in China. Vet. Microbiol. 230, 56–61 (2019).

4. Silva-Rocha, R. et al. The Standard European Vector Architecture (SEVA): a coherent platform for the analysis and deployment of complex prokaryotic phenotypes. Nucleic Acids Res. 41, (2013).

